# Primary cilia regulate GLP-1 signaling in pancreatic β cells

**DOI:** 10.64898/2026.02.22.707280

**Authors:** Isabella Melena, Jeong Hun Jo, Shannon E. Townsend, Samantha Adamson DiGruccio, Xinhang Dong, Lifei Zhu, Jonathan Campbell, Jing W. Hughes

**Affiliations:** Department of Medicine, Division of Endocrinology, Metabolism & Lipid Research, Washington University School of Medicine, St. Louis, MO 63110, USA; Department of Medicine, Yale University School of Medicine, New Haven, CT 06510, USA; Department of Medicine, Duke University School of Medicine, Durham, NC 27710, USA

## Abstract

Glucagon-like peptide-1 receptor agonists (GLP-1RAs) are mainstay therapies for diabetes and obesity, acting in part by enhancing glucose-dependent insulin secretion. While the primary cilium is a known signaling compartment for certain G-protein coupled receptors (GPCRs), its role in the β-cell response to incretins remains undefined. Here, we show that primary cilia are essential for GLP-1R signaling. Loss of β-cell cilia in mouse and human islets severely impaired GLP-1-potentiated insulin secretion, an effect preceded by blunted whole-cell cAMP and Ca²⁺ responses. Immunofluorescence and immunogold scanning electron microscopy revealed endogenous GLP-1R localized to the primary cilium. Critically, disrupting ciliary GPCR trafficking via Tulp3 knockdown – while preserving cilia structure – recapitulated the signaling and secretory deficits, demonstrating a specific requirement for the ciliary receptor pool. These findings establish the primary cilium as a non-redundant signaling compartment for GLP-1R and uncover a new layer of subcellular organization in incretin action in β cells.

## Introduction

Glucagon-like peptide-1 receptor agonists (GLP-1RAs) are cornerstone therapies in diabetes and obesity, partly due to their ability to enhance β-cell glucose-dependent insulin secretion. The GLP-1 receptor (GLP-1R) is a G-protein coupled receptor (GPCR) that activates both G_q_ and G_s_ pathways, leading to elevation of cytosolic Ca^2+^ and cAMP that collectively potentiate insulin exocytosis^1–5^. However, these second messengers are ubiquitous intracellular signals, and it remains unclear how GLP-1R activation generates a specific response to stimulate insulin release. Although GLP-1R is known to operate in discrete subcellular domains including the plasma membrane where it initiates signaling and endosomes where it can sustain cAMP production^6–8^, additional undefined spatial mechanisms likely contribute to its signaling output. The compartmentalization of receptors and their effector molecules is increasingly recognized as a critical determinant of signaling specificity and amplitude that enable precise cellular control. Elucidating these microdomains for GLP-1R could therefore clarify the efficacy of GLP-1RAs and help explain the variation in individual treatment responses observed clinically^9,10^.

Emerging evidence identifies primary cilia as privileged platforms for metabolic GPCR signaling. These microtubule-based organelles, present on most mammalian cells, concentrate receptors and effector molecules within a diffusion-restricted compartment, creating a unique biochemical environment that amplifies signaling potency^11,12^. This location bias allows ciliary pools of GPCRs to influence cellular dynamics in ways distinct from their actions on the plasma membrane. This principle has been demonstrated in neurons and kidney cells, where ciliary GPCRs including the melanin-concentrating hormone receptor 1 (MCHR1), melanocortin 4 receptor (MC4R), leptin receptor (LepR), prostaglandin receptor (EP4), and polycystin channels (PC1 and 2) drive whole-cell signaling responses critical for cellular metabolism and homeostasis^13–18^. In pancreatic islets, several paracrine hormone GPCRs, such as somatostatin receptor (SSTR3/5) and GABA_B1_ receptor, are strongly enriched in the primary cilium, where they modulate local and cytosolic cAMP and Ca²⁺ dynamics to influence insulin secretion^19–21^. Ciliary GPCR localization depends on specific trafficking machinery, particularly the TULP-IFTA complex which mediates the entry of membrane proteins like SSTR3 into the ciliary compartment^22,23^. Furthermore, genetic ablation of β-cell cilia disrupts glucose- and paracrine-regulated insulin release and promotes diabetes *in vivo*, underscoring the physiological relevance of ciliary signaling to metabolic health^24,25^.

Given the cilium’s established role as a metabolic signaling hub, we hypothesized that primary cilia are required for the β-cell response to GLP-1. To test this, we first examined the functional consequences of cilia loss-of-function. Using mouse islets lacking β-cell cilia and human islets following knockdown of IFT88 (a core component of the intraflagellar transport machinery required for cilia assembly), we found that GLP-1-potentiated insulin secretion was significantly impaired. This secretory defect was preceded by blunted whole-cell cAMP and Ca²⁺ responses, indicating an upstream signaling failure. To determine whether this failure depends on the subcellular localization of GLP-1R, we performed high-resolution imaging of endogenous receptor distribution, revealing a distinct pool of GLP-1R within the primary cilium of pancreatic islet cells, suggesting the cilium may serve as a required signaling compartment. To functionally test this model, we disrupted ciliary GPCR trafficking via Tulp3 knockdown, which recapitulated the signaling and secretory deficits without destroying cilia structure, showing that proper ciliary localization of GLP-1R itself is essential. Together, these data establish that the primary cilium functions as a non-redundant signaling compartment for GLP-1R and that the ciliary receptor pool is necessary for generating whole-cell incretin responses.

## Results

### β-cell primary cilia are required for GLP-1 augmented insulin secretion

Activation of the GLP-1R potently enhances insulin secretion selectively under high glucose conditions, a mechanism critical to its safety as an anti-diabetic agent^2,26,27^. Building on evidence that the GLP-1R mediates intra-islet paracrine signaling to regulate secretion^1,4^, and our previous work showing that β-cell primary cilia integrate glucose and paracrine signals^19,25^, we investigated whether cilia are required for GLP-1R action. We first performed islet perifusion using β-cell-specific cilia knockout (βCKO) and wild-type (WT) mice (**Figure 1A**). While WT islets robustly increased insulin secretion upon stimulation with 16 mM glucose and subsequent 20 nM liraglutide (a GLP-1 receptor agonist), βCKO islets exhibited a significantly blunted secretory response to both stimuli (**Figure 1A-C**). This impairment was specific to receptor-mediated pathways, as depolarization-induced secretion with 30 mM KCl remained intact (**Figure 1D**), indicating an unaffected exocytotic machinery. To confirm that the defect was not due to altered receptor expression, we labeled whole islets with the fluorescent GLP-1R antagonist LUXendin645 and found comparable fluorescence intensity between WT and βCKO islets (**Figure S1**). This indicates that total islet GLP-1R protein levels are unchanged by cilia loss. Quantitative analysis of the perifusion data revealed that βCKO islets secreted ∼50% less insulin during the liraglutide phase (AUC, minutes 30-50) than WT controls (**Figure 1B**), and total secreted insulin over the entire time course was significantly reduced (**Figure 1C**). Together, these data demonstrate that primary cilia are required for the full secretory response to GLP-1R activation, independent of changes in total receptor abundance.

**Figure 1:**
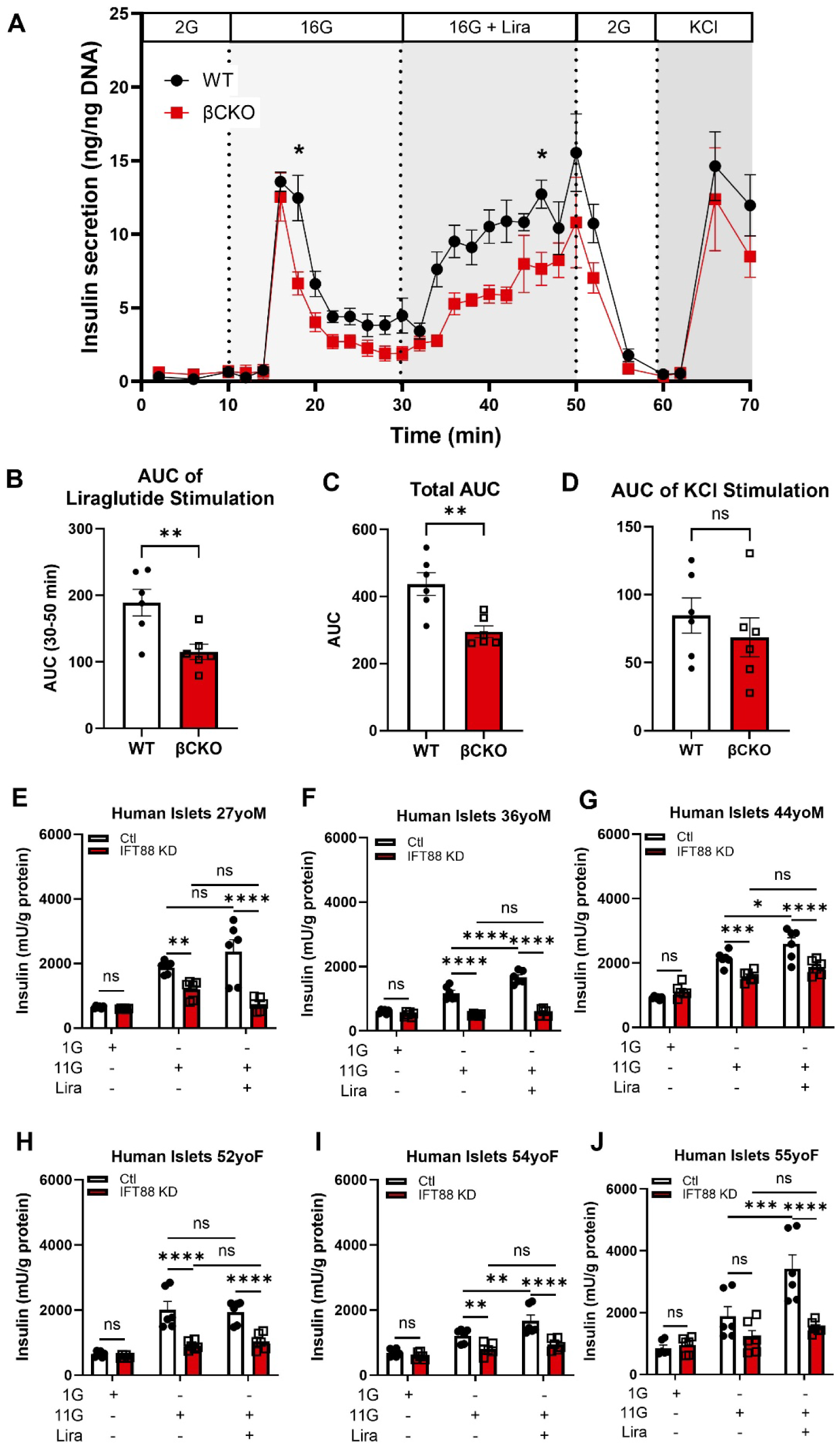
Primary cilia are required for GLP-1-potentiated insulin secretion. (**A**) Dynamic insulin secretion from perifused mouse islets. Wild-type (WT, black) and β-cell-specific cilia knockout (βCKO, red) islets were sequentially exposed to 2 mM glucose (2G), 16 mM glucose (16G), 20 nM liraglutide (Lira), and 30 mM KCl as indicated. N = 6 replicates of 50 islets per genotype from 5 mice. *p<0.05, two-way ANOVA with Sidak’s multiple comparisons test. (**B–D**) Quantitative analysis of secretion traces. (**B**) Area under the curve (AUC) during liraglutide stimulation (minutes 30–50). (**C**) Total AUC of the entire perifusion (minutes 0-70). (**D**) AUC during KCl depolarization (minutes 60–70). βCKO islets secreted significantly less insulin in response to liraglutide and overall but showed no defect in KCl-induced secretion. Data are mean ± SEM. **p<0.01; ns, not significant, unpaired student’s t-test. (**E–J**) *IFT88* knockdown impairs GLP-1-augmented secretion in human islets. Static glucose-stimulated insulin secretion (GSIS) assays from islets of male (**E–G**) and female (**H–J**) donors. Islets transduced with control (Ctl, white) or *IFT88*-targeting shRNA (IFT88 KD, red) were incubated at 1 mM glucose (1G), 11 mM glucose (11G), and 11G + 100 nM liraglutide (Lira). Donor ages are indicated. IFT88 KD significantly reduced liraglutide-potentiated insulin secretion. Data are mean ± SEM of triplicate samples per donor. *p<0.05, **p<0.01, ***p<0.001, ****p<0.0001; ns, not significant; one-way ANOVA with Tukey’s multiple comparisons test.

To extend these findings to human islets, we knocked down the essential ciliary gene *IFT88* using adenoviral shRNA (IFT88 KD), achieving a 50-70% reduction in whole-cell IFT88 expression and reduction in mean cilia length (**Figure S2**). An expanded distribution of cilia length was also observed in IFT88 knockdown, consistent with bidirectional length alterations reported for IFT deletions^28–31^. Functionally, IFT88 KD significantly impaired glucose-stimulated insulin secretion in five of six human donors tested (**Figure 1E-I**), mirroring the mouse phenotype. Notably, in donors where liraglutide potentiated secretion, this potentiation was markedly attenuated in IFT88 KD islets compared to controls (**Figure 1E-J**). This finding provides functional evidence that disrupted cilia impairs the local intra-islet action of GLP-1R in human β cells. Consistent with the known heterogeneity of human islet function and incretin responsiveness, baseline liraglutide potentiation varied considerably across a broader cohort of 20 control (non-KD) donors, with significant potentiation observed in only a subset (**Figure S3, Table S1**).

### Cilia potentiate GLP-1-stimulated cAMP and Ca^2+^ dynamics

#### cAMP signaling

To dissect the mechanism underlying the secretory defect, we first measured GLP-1-evoked cAMP dynamics in live mouse islets. This was done using the FRET-based sensor Epac-SH187 (**Figure 2A**), an established approach for monitoring cAMP in intact islets where the signal is predominantly β-cell-derived^32^. Stimulation with 100 nM liraglutide at 11 mM glucose elicited a robust rise in cAMP in WT islets, which was significantly blunted in βCKO islets (**Figure 2B**). This indicates that primary cilia are necessary for a normal GLP-1R-mediated cAMP response. We also observed that the cAMP deficit in βCKO islets extended to conditions of direct adenylyl cyclase activation with forskolin (FSK) and with combined FSK and IBMX (**Figure 2C-D**). While this indicates a more general impairment in cAMP production or accumulation in cilia-deficient β cells, the cAMP defect under GLP-1R-specific stimulation is consistent with a direct requirement for cilia in incretin receptor G_s_ signaling.

**Figure 2:**
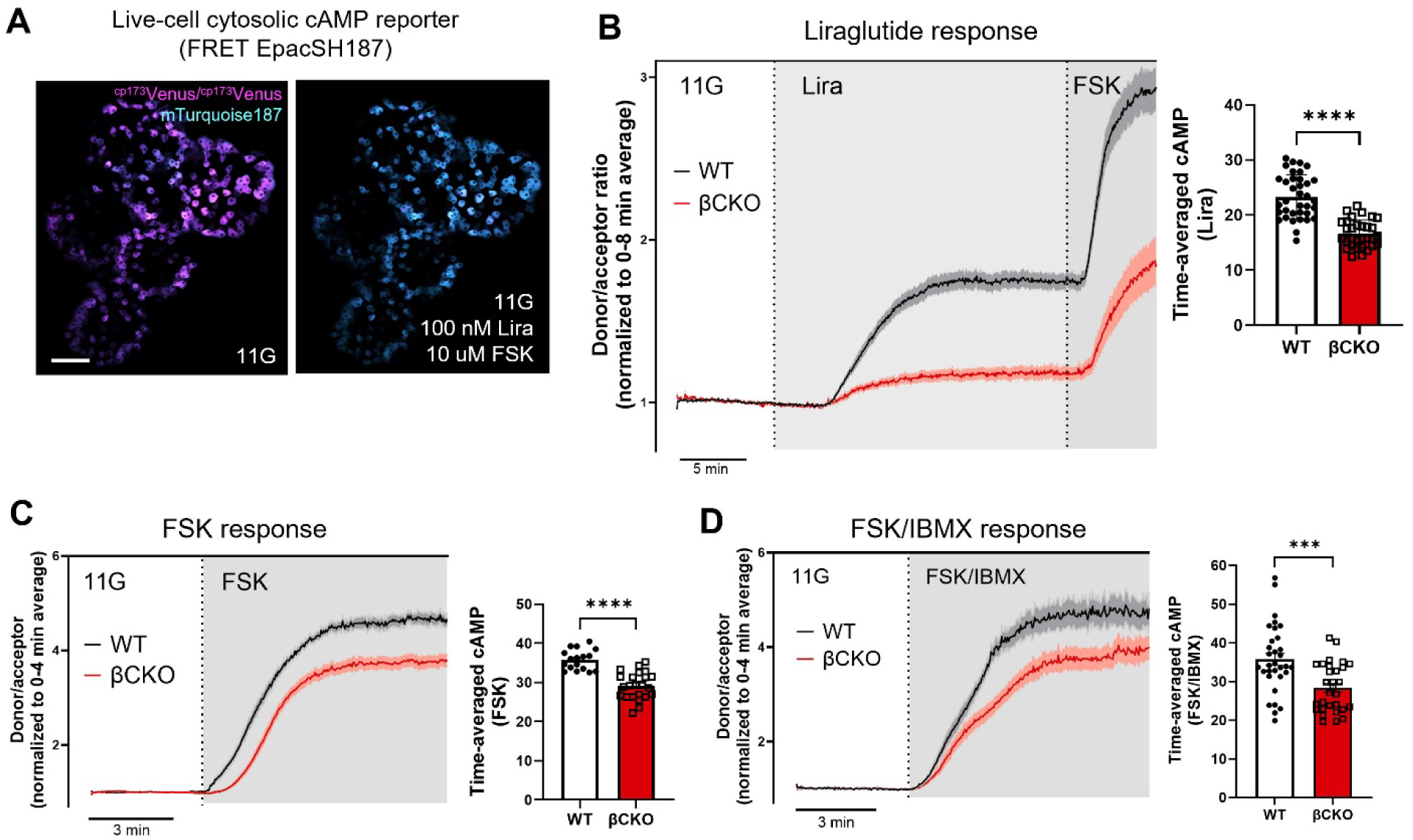
Cilia loss impairs cAMP generation in response to GLP-1 and direct adenylyl cyclase activation. (**A**) Representative FRET images of WT islets expressing the EpacSH187 cAMP sensor before (left) and after (right) stimulation with 100 nM Liraglutide (Lira) and 10 µM forskolin (FSK). Pseudocolors show the acceptor (^cp173^Venus, magenta) and donor (mTurquoise187, cyan) channels. Scale bar = 20 µm. (**B**) Average FRET ratio traces (mean ± SE) from WT (black) and βCKO (red) islets stimulated with 11 mM glucose (11G), with sequential addition of 100 nM Lira and 10 µM FSK as indicated. Quantification shows time-averaged cAMP response during Lira stimulation (minutes 8-23): βCKO islets generated significantly less cAMP than WT. n= 36 WT, 34 βCKO islets from 4 mice per genotype. (**C**) Average FRET ratio traces (mean ± SE) from WT and βCKO islets in 11G, stimulated with 10 µM FSK. Quantification shows total time-averaged cAMP response where βCKO islets exhibited reduced response. N= 32 WT, 30 βCKO islets from 3 mice per genotype. (**D**) Average FRET ratio traces (mean ± SE) from WT and βCKO islets in 11G, stimulated with 10 µM FSK and 100 µM IBMX. Quantification showing βCKO islets generated less cAMP than WT. n = 25 WT, 23 βCKO islets from 3 mice per genotype. ***p<0.001, ****p<0.0001; student’s unpaired t-test.

#### Ca^2+^ signaling

We next analyzed GLP-1-modulated Ca^2+^ dynamics in β-cell GCaMP6f reporter mice. As reported^33^, liraglutide increased β-cell Ca^2+^ oscillation frequency under stimulatory glucose conditions. In our study, βCKO islets exhibited a delayed and diminished initial Ca^2+^ response to 9 mM glucose (**Figure 3A**), consistent with our previous findings^19^. The augmentation of second-phase oscillations by 20 nM liraglutide was blunted in βCKO islets, resulting in a significantly lower total Ca^2+^ flux (AUC, **Figure 3B-D**). Waveform analysis showed that liraglutide-evoked oscillations in βCKO islets had reduced amplitude, a shorter period, and decreased duration of both active and silent phases compared to WT, indicating an accelerated but weakened oscillatory cycle (**Figure 3E-H**). Similarly, exendin-4 (Ex4) generated defective Ca^2+^ signals in βCKO islets, characterized by a shortened first peak and altered steady-state oscillation kinetics (**Figure S3**). Together, these results demonstrate that cilia loss prevents the normal amplification of Ca^2+^ oscillations by GLP-1R agonists, resulting in a reduced integrated Ca^2+^ signal. In sum, these findings indicate that primary cilia help transduce GLP-1 signals into the canonical cAMP and Ca^2+^ second-messenger pathways that drive insulin secretion.

**Figure 3:**
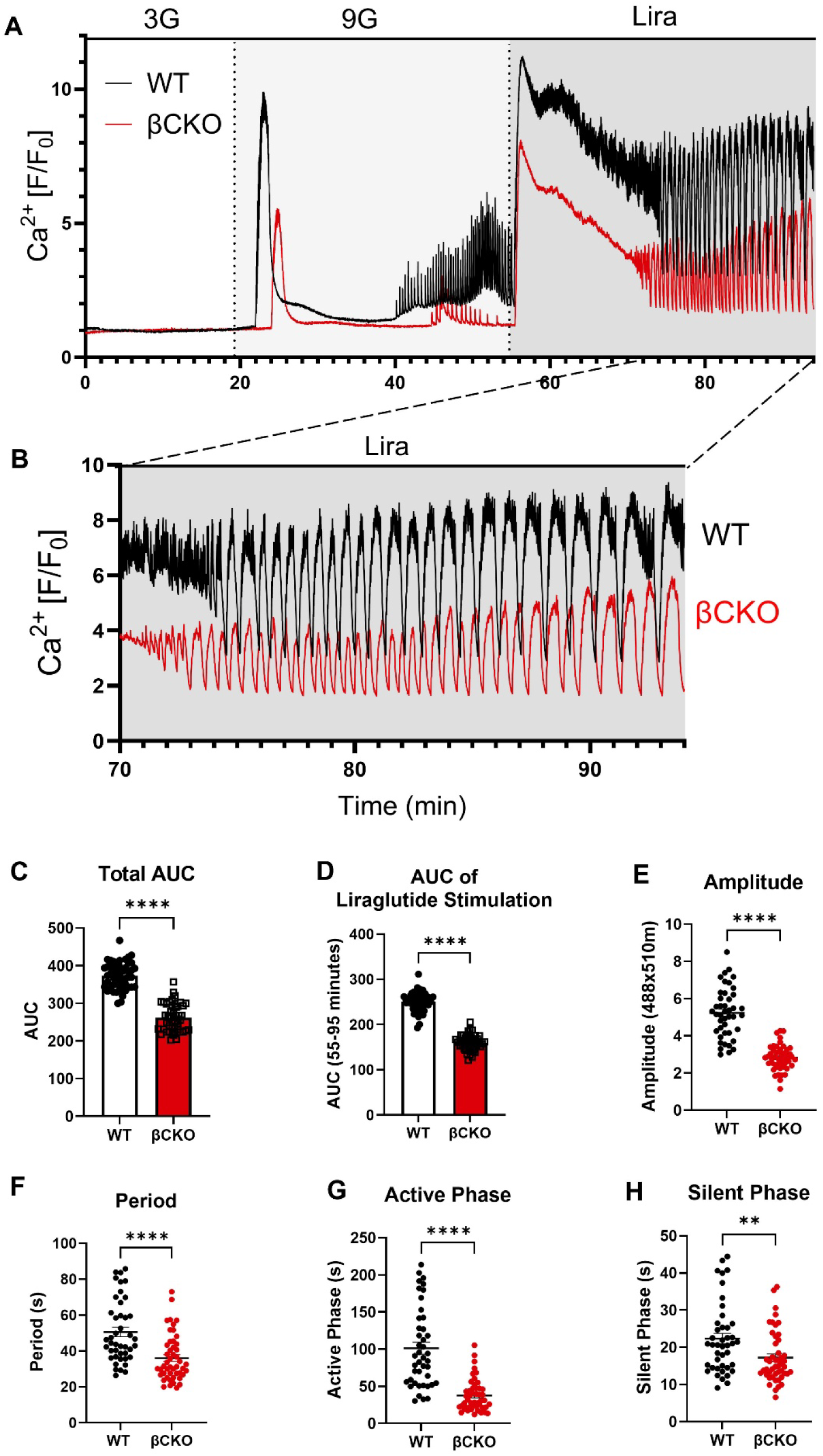
Cilia loss impairs GLP-1-augmented Ca^2+^ dynamics. (**A**) Representative single-islet Ca²⁺ traces from WT (black) and βCKO (red) islets, showing normalized GcaMP6f fluorescence over time. Islets were stimulated with 9 mM glucose (9G), followed by addition of 100 nM liraglutide (Lira) at the time indicated. (**B**) Representative single-islet Ca²⁺ traces from WT and βCKO islets of oscillations with Lira, expanded from (A) as indicated by the dashed lines. (**C**) Total area under the curve of traces shown in A between WT and βCKO islets. (**D**) Area under the curve of Lira stimulation from 55-95 minutes of traces in (A) between WT and βCKO islets. (**E-H**) Waveform analysis of Ca² oscillations during liraglutide stimulation (70-95 min period in **B**). I Amplitude, (**F**) period, (**G**) active phase, and (**H**) silent phase. N= 49 WT and 49 βCKO islets, n=3 mice each. Statistical significance was assessed by student’s unpaired t-test: **p<0.01, ***p<0.001, ****p<0.0001.

### A specific pool of GLP-1R localizes to the primary cilium

Primary cilia concentrate specific GPCRs critical for their signaling function. To determine if GLP-1R is among these receptors, we performed high-resolution localization studies on mouse islet cells. Using a recently validated monoclonal antibody (DSHB Mab7F38)^34^, paired with high-performance secondary dyes (Abberior STAR), we detected endogenous GLP-1R on primary cilia by confocal microscopy (Figure 4A). Immunogold scanning electron microscopy (1mmune-SEM) confirmed GLP-1R localization within the ciliary shaft at ultrastructural resolution (**Figure 4B**). Specificity of the immunostaining was validated using islets from β-cell-specific *Glp1R/GcgR* double knockout mice; these controls showed significantly ablated GLP-1R staining signal, including its absence from ciliary structures (**Figure 4C-D**). Importantly, GLP-1R protein and mRNA levels were largely unchanged in βCKO islets (**Figure 4E-F**), indicating that cilia loss impairs signaling downstream of the receptor rather than altering its expression or global distribution.

**Figure 4:**
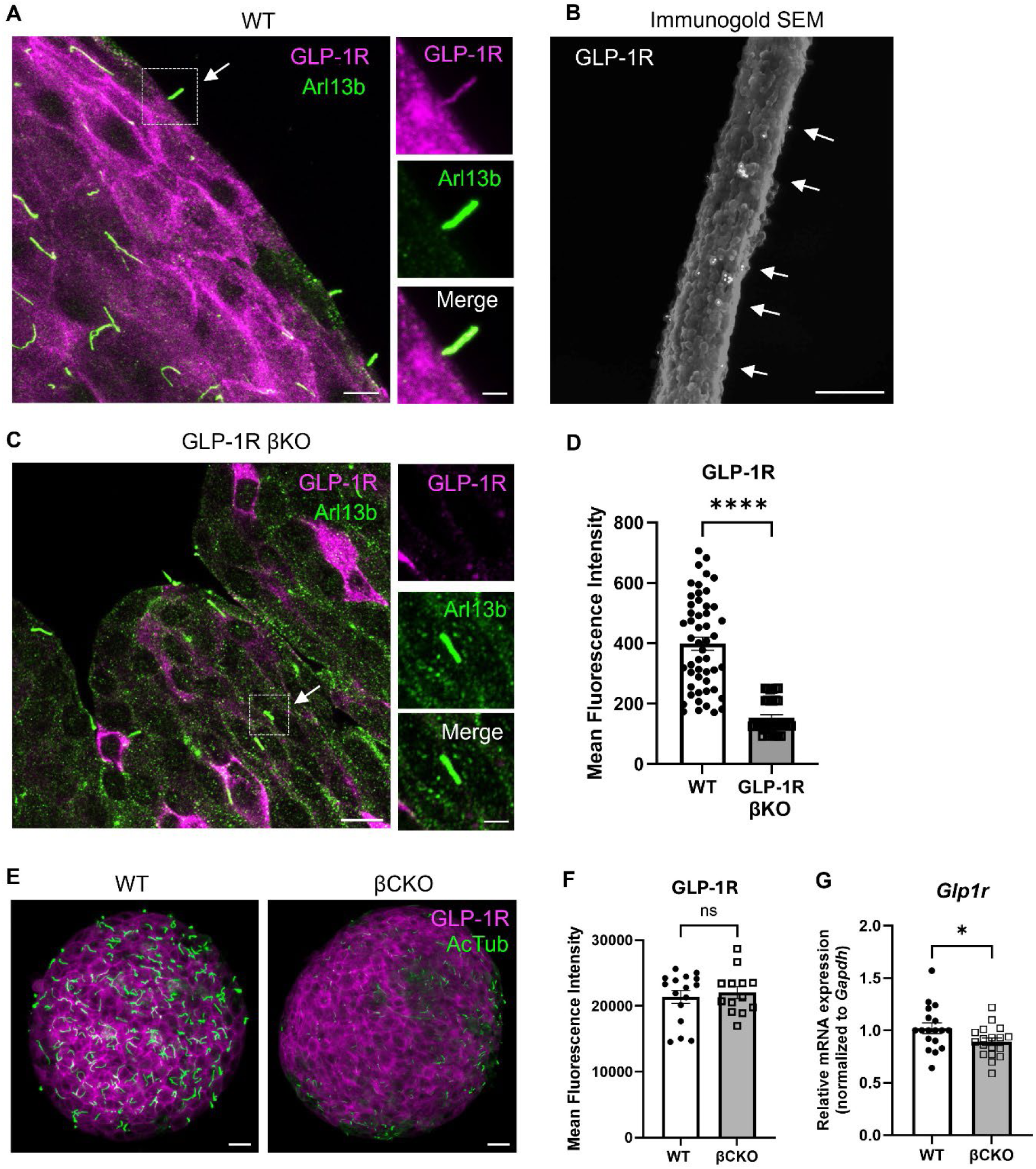
GLP-1R localizes to islet primary cilia. (**A**) Representative confocal image of a WT islet immunostained for GLP-1R (magenta) and the ciliary marker Arl13B (cilia, green); scale bar = 10 µm. Inset depicts single-channel images of the cilium confirming GLP-1R colocalization with Arl13b; scale bar = 3 µm. (**B**) Immunogold scanning electron microscopy of a single cilium. Gold particles (white arrows) indicate GLP-1R labeling. Scale bar = 300 nm. (**C**) Representative confocal image of a GLP-1R β-cell knockout (KO (GLP-1R βKO) islet stained as in (A), showing loss of GLP-1R signal in the cilium (dashed box). Scale bar =10 µm. (**D**) Quantitation of normalized GLP-1R fluorescence intensity in single optical sections of WT and GLP-1R βKO islets. N = 8 WT, 10 GLP-1R βKO islets. I Representative whole-islet images of WT and βCKO islets stained for GLP-1R (magenta) and cilia marker acetylated tubulin (AcTub, green), showing reduced cilia in βCKO islets without a corresponding change in total GLP-1R signal. (**F**) Quantitation of mean GLP-1R intensity from 5 µm z-projections from whole islets, showing no difference between WT and βCKO. N = 19 WT, 20 βCKO islets. (**G**) Relative *Glp1r* mRNA expression in WT and βCKO islets. Expression was significantly reduced in βCKO islets. Data are from triplicate samples across two independent experiments. Data are mean ±SEM. Statistical significance was assessed by student’s unpaired t-test: *p<0.05, **p<0.01, ***p<0.001, ****p<0.0001; ns not significant.

### Disrupting ciliary GPCR trafficking blunts GLP-1 responsiveness

To directly test the functional role of the ciliary GLP-1R pool, we disrupted its trafficking to the cilium without ablating the structure itself. We targeted TULP3, an intraflagellar transport (IFT) adapter protein that shuttles a specific subset of GPCRs (including SSTR3) into the ciliary compartment^23,35^. In WT islets, β-cell-targeted knockdown of *Tulp3* (Tulp3 KD) did not alter overall GLP-1R expression, as assessed by immunostaining and mRNA levels (**Figure 5A-D**), nor did it disrupt ciliary architecture (**Figure 5G**). However, it significantly reduced GLP-1R fluorescence intensity specifically at the cilium (**Figure 5E, F**), confirming a selective defect in ciliary receptor localization. Functionally, Tulp3 KD islets exhibited a signaling defect, with a significantly blunted rise in liraglutide-augmented cAMP (**Figure 5H, I**). This impairment translated to secretion, where liraglutide-potentiated insulin output was markedly reduced, while the secretory response to glucose alone remained intact (**Figure 5J**). Together, these data demonstrate that proper ciliary localization of GLP-1R, mediated by TULP3-dependent trafficking, is necessary for its full incretin activity.

**Figure 5:**
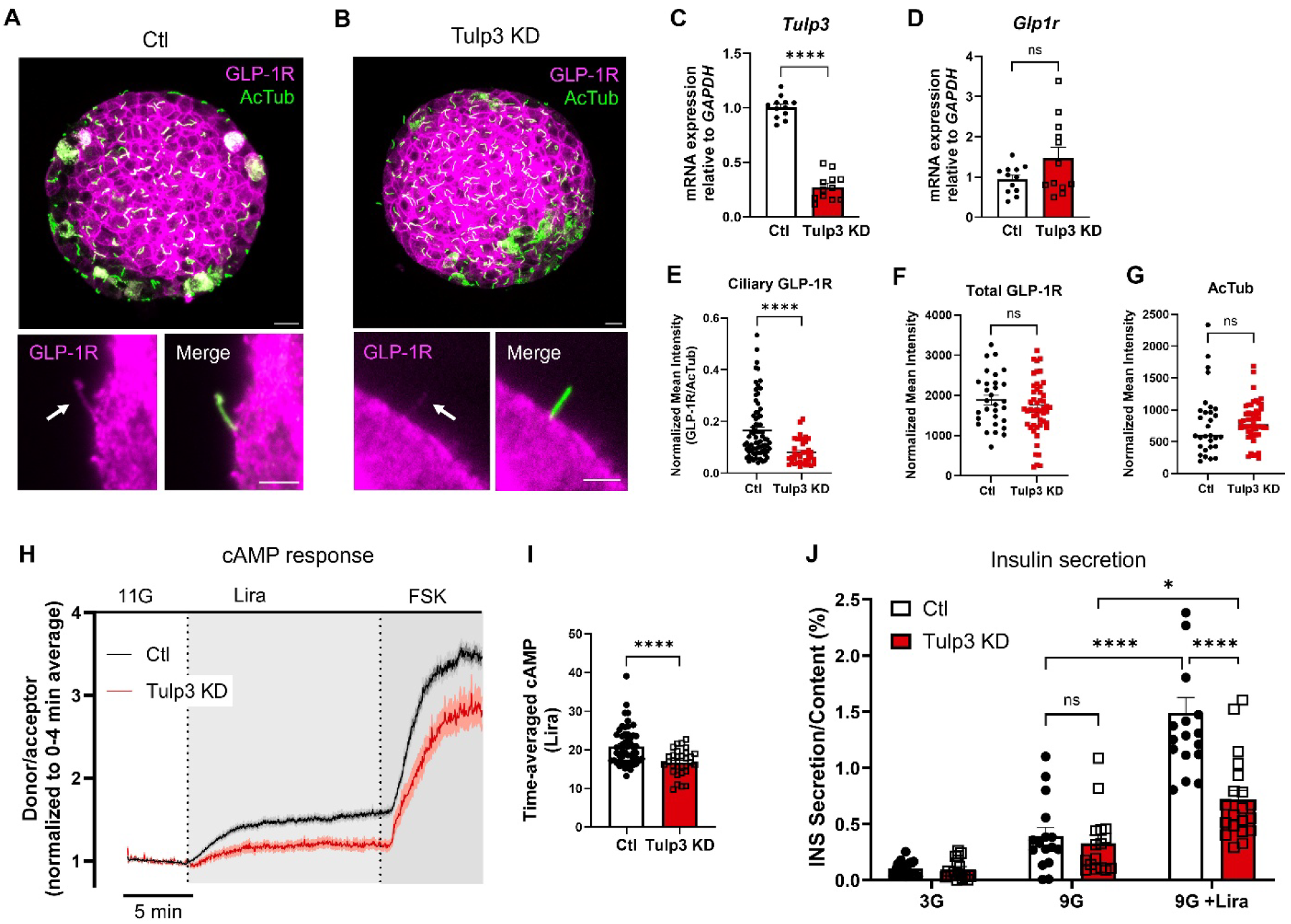
Tulp3 is required for ciliary localization of GLP-1R and for GLP-1-mediated signaling and secretion. (**A-B**) Localization of GLP-1R (magenta) and the ciliary marker acetylated tubulin (AcTub, green) in control (Ctl) and *Tulp3* knockdown (Tulp3 KD) islets. The main images show whole islets; insets display magnified single cilia in individual channels. (**C-D**) Relative mRNA expression of *Tulp3* I and *Glp1r* (**D**) in control and Tulp3 KD islets. Knockdown selectively reduced *Tulp3*, while *Glp1r* levels were unchanged. Data are from triplicate samples across four independent experiments. I Quantification of cilia-specific GLP-1R fluorescence intensity, showing reduced receptor localization in Tulp3 KD cilia. n = 74 control and 38 Tulp3 KD cilia from 8-9 mice. (**F**) Whole-islet GLP-1R intensity from z-projections, showing no difference between control and Tulp3 KD. This finding is consistent with the unchanged surface labeling by LUXendin645 (**Figure S1**). N= 30 control and 50 Tulp3 KD islets. (**G**) Quantification of total cilia length and number per islet, based on AcTub staining. N= 30 control and 50 Tulp3 KD islets (**H**) Average FRET ratio traces (mean ± SE) from control (black) and Tulp3 KD (red) islets expressing the EpacSH187 cAMP sensor. Islets were placed in 11 mM glucose (11G) followed by sequential addition of 100 nM Lira and 10 µM forskolin (FSK). (**I**) Quantification of the time-averaged cAMP response during liraglutide stimulation (minutes 4-20). Tulp3 KD significantly blunted the cAMP rise. N=53 control and 31 Tulp3 KD islets from 9 mice. (**J**) Insulin secretion from control and Tulp3 KD islets at 3 mM glucose (3G), 9 mM glucose (9G), and 9G + 100 nM liraglutide (Lira). Tulp3 KD impaired liraglutide-potentiated secretion. N= 18 replicates of 5 islets per condition across three experiments. Data are mean ±SEM. Statistical significance was assessed by student’s unpaired t-test: *p<0.05, **p<0.01, ***p<0.001, ****p<0.0001; ns, not significant.

## Discussion

This work provides evidence that the primary cilium constitutes a functional, non-redundant compartment for GLP-1R signaling in pancreatic β cells. While GLP-1R signaling has been characterized at the plasma membrane and in endosomes, our results indicate that a ciliary-localized pool of the receptor is required to generate the full insulin secretory response to GLP-1 receptor agonists (GLP-1Ras).

Our conclusion is supported by several independent lines of evidence. High-resolution imaging of endogenous GLP-1R protein confirmed its localization to the primary cilium of mouse islet cells, a result that was independently confirmed by another laboratory (Huising Lab, personal communication). Genetic disruption of cilia in mouse (βCKO) and human islets (IFT88 knockdown) reduced GLP-1RA-potentiated insulin secretion by approximately 50%, a defect accompanied by blunted whole-cell cAMP and Ca²⁺ responses. Importantly, selectively disrupting ciliary GPCR trafficking through Tulp3 knockdown, which diminishes ciliary GLP-1R localization without ablating the cilium, recapitulated the same signaling and secretion defects, suggesting that the physical presence of the receptor within the cilium is a key determinant of signaling output. The phenotype was specific to receptor-potentiated secretion, as the insulin response to direct membrane depolarization with KCl remained intact, indicating that cilia are not universally required for exocytosis but participate specifically in the GLP-1R-mediated amplification pathway.

The observed defects in cAMP and Ca²⁺ signaling suggest that the primary cilium serves as an upstream node that influences whole-cell signaling and determines the robustness of the β-cell response to incretin stimulation. The reduction in cAMP accumulation in βCKO islets, even upon combined FSK/IBMX treatment, indicates that cilia may contribute to the overall capacity to generate and maintain this second messenger. In parallel, the cilia-dependent Ca²⁺ changes to glucose and GLP-1RA suggest a perturbation of the β-cell oscillatory machinery, in which both phases of the cycle are reduced in the absence of the cilium, thus accelerating oscillatory frequency and weakening the amplitude. Speculatively, this could reflect a shortened refractory period (e.g. enhanced Ca²⁺ extrusion or K⁺ channel kinetics) or a reduced driving force (e.g. impaired metabolic coupling or diminished Ca²⁺-induced Ca²⁺ release), shifting the system into a faster but less robust rhythm. While direct electrophysiological evidence is needed to distinguish these models, the data collectively suggest that the cilium helps set the gain and timing of the Ca²⁺ oscillator, which in turn modulates the secretory response to incretin input.

Our study raises several key mechanistic questions. First, the diffusion-restricted volume of the cilium makes it unlikely that ciliary-derived second messengers (Ca²⁺, cAMP) themselves directly contribute meaningfully to the global cytosolic pool. While ciliary cAMP likely does not measurably impact whole-cell cAMP levels, our functional data demonstrate that the cilium nevertheless exerts a significant influence on global cAMP dynamics. This effect is not explained by changes in plasma membrane GLP-1R expression, as LUXendin labeling confirmed comparable surface receptor levels in WT and βCKO islets (**Figure S1**). Instead, the cilium may act as a regulatory initiator or amplifier, possibly through localized effectors such as PKA, local metabolite generation, or physical coupling of signaling microdomains. Cilia could also modulate the broader cAMP signaling machinery, such as by regulating the expression or activity of ACs and PDEs, thereby influencing whole-cell cAMP generation or maintenance independently of receptor abundance. Our pilot immunostaining of pan-ACs revealed no overt differences between WT and cilia KO cells, but these results do not exclude functional changes at the level of subcellular localization or enzyme activity. Addressing these possibilities would require going beyond fixed-cell staining involving more sophisticated tools such as live-cell biosensors and spatially resolved biochemical approaches.

Second, while Tulp3 knockdown established that ciliary GLP-1R localization is necessary for full β-cell activation, direct measurement of compartment-specific second-messenger dynamics is still needed to determine whether the ciliary pool engages unique downstream effectors. The nature of the outgoing ciliary signal also remains to be determined. Several speculative and non-mutually exclusive mechanisms could explain the observed functional connection between cilia and cell body: the cilium could serve as a signaling scaffold that organizes and amplifies downstream effectors, e.g. PKA or EPAC, at the ciliary base, which then propagates signals into the cytosol. Cilia could organize AKAP-anchored signaling complexes that tether PKA to key effector sites like K_ATP_ channels near the plasma membrane, coupling G-protein signaling to β-cell excitation^15,36^. Additionally, ciliary signaling could influence cytosolic cAMP dynamics by regulating PDE activity at the ciliary base, or by modulating cytosolic Ca^2+^ (**Figures 3, S4**) to affect cAMP production via calcium-sensitive ACs. Future work should also investigate post-activation mechanisms, including potential β-arrestin recruitment and BBSome-mediated export^37^, to understand local actions of GLP-1R within the cilium and the regulation of signal duration and receptor recycling. Although Tulp3 traffics multiple ciliary cargos, our finding that Tulp3 KD impairs GLP-1 action without altering total GLP-1R expression demonstrates a specific role for ciliary targeting in this pathway. Identifying GLP-1R ciliary-targeting sequences and testing cilia-targeting mutants will help distinguish receptor-specific effects from broader trafficking mechanisms. Finally, our functional data rely primarily on liraglutide; further study is required to determine whether ciliary dependence extends to other GLP-1Ras or to endogenous GLP-1.

From a translational perspective, the donor-to-donor variability in human islet responses to liraglutide observed here and elsewhere highlights the challenge of incretin research and the clinical challenge of incretin-based therapies. Our findings raise the possibility that inter-individual differences in ciliary structure, protein content, or trafficking efficiency may contribute to this heterogeneity. Future studies examining whether ciliary GLP-1R signaling is compromised in diabetic islets, and whether this contributes to therapeutic non-responsiveness, might provide insights into personalized diabetes management using incretin mimetics.

## Materials and methods

### Mice

All mouse experiments were approved by the Washington University Institutional Animal Care and Use Committee. Male and female mice aged 8–14 weeks were used for all experiments. Wild-type Ins1-Cre and IFT88^fl/fl^ mice, along with β-cell cilia knockout mice (βCKO; Ins1-Cre x IFT88^fl/fl^) as previously described^25^, were used for perifusion hormone secretion assays and cAMP imaging. Ins1-Cre GcaMP6f and βCKO GcaMP6f mice as previously described^19^ were used for Ca^2+^ imaging. GLP-1R βKO (GLP-1R^f/f^ MIP-CreERT2^+/+^) and wild-type (GLP-1R^f/f^ MIP-CreERT2^-/-^) were contributed by the Campbell Lab following *in vivo* tamoxifen treatment for KO induction^1,38^. Healthy non-diabetic human islets were obtained from the Integrated Islet Distribution Program (IIDP). Donor information is provided in **Table S1**.

### Pancreatic islet isolation and culture

Murine pancreatic islets were isolated from 8-14 week old mice through collagenase P digestion (Thermo Fisher Scientific). Following digestion, islets were either hand-picked under stereomicroscopic guidance or purified by histopaque as previously described^25,39,40^, then subsequently maintained overnight in RPMI 1640 medium supplemented with 10% FBS and 1% penicillin/streptomycin at 37°C (5% CO_2_) prior to experiments.

### Immunohistochemistry

Isolated islets were fixed with 4% formaldehyde (FA) for 15 min at room temperature and rinsed three times with PBS. Samples were blocked and permeabilized at room temperature for 1 hour in CAS-Block Histochemical Reagent (Thermo Fisher Scientific) with 0.3% TritonX-100, then incubated with primary antibodies for 24-48 h at 4°C. After primary incubation, islets were washed extensively with 5 x 30-min exchanges in PBST (PBS with 0.3% Triton X-100), then incubated with species-appropriate secondary antibodies overnight at 4°C or 1 h at room temperature, followed by 3-5 PBST washes. Islets were mounted in aqueous mounting medium and stored at 4°C protected from light until imaging.

Primary antibodies included: GLP-1R (DSHB, Mab7F38, 1:50), ARL13B (Proteintech, 30332-1-AP, 1:400), AcTub (Proteintech, 66200-1-Ig; Sigma-Aldrich, T7451, 1:400), Tulp3 (Proteintech, 13637-1-AP, 1:200). Secondary antibodies included Abberior red and far-red STED dyes (STAR-orange, STAR-red; 1:200) and Alexa Fluor goat anti-mouse, anti-rabbit, and anti-guinea pig (Invitrogen, 1:500). Live-cell GLP-1R labeling was performed using LUXendin645 (Celtarys, 100 nM) per published protocol^41^.

Confocal imaging was performed using a Nikon AXR microscope equipped with Nikon spatial array confocal (NSPARC) detector (Nikon, Tokyo, Japan). Images were acquired using Plan Apochromat 60x and 100× silicone oil immersion objectives (NA 1.35). All experiments used the Galvano scanner with the following parameters: 2 µs dwell time, 2x scan averaging, and 2048 x 16-pixel resolution. Laser power and z-stack settings were optimized for each channel to maintain consistent signal intensity throughout the imaging volume while using approximately 30% of the detector’s dynamic range.

### Immunogold electron microscopy

Adherent mouse islets were rinsed in PBS, quenched with 50 mM glycine, and blocked for 30 min in PBS containing 1% BSA and 5% normal donkey serum. Samples were incubated with primary antibody (mouse anti-GLP-1R, DSHB Mab7F38) diluted in blocking solution for 24 h at 4°C followed by 5 h at room temperature; negative controls omitted the primary antibody. After five 5-min washes in blocking solution, islets were labeled for 1.5 hr with 18-nm gold-conjugated secondary antibody (donkey anti-mouse/rabbit, 1:20) in blocking buffer. Following washes in blocking solution and PBS, samples were fixed in fresh 2% glutaraldehyde (15 min), stained with 0.5% osmium tetroxide (20 min on ice), and rinsed in Milli-Q water. Dehydration was performed through a graded ethanol series (10–100%), after which islets were critical-point dried, coated with a 12-nm carbon layer, and stored in a desiccator. Imaging was performed on a Helios SEM at 5 kV and 0.1 nA.

### Static secretion assay

Islets were equilibrated in Krebs-Ringer bicarbonate HEPES (KRBH) buffer (128.8 mM NaCl, 4.8 mM KCl, 1.2 mM KH_2_PO_4_, 1.2 mM MgSO_4_, 2.5 mM CaCl_2_, 20 mM HEPES, 5 mM NaHCO_3_, H_2_O, and 0.1% BSA [pH 7.4]) at 2.8 mM glucose for 45 min at 37°C. Following equilibration, islets (5-10 per tube) were stimulated for 1 h at 37°C with either low (1-3) or high (9-11) mmol/L glucose conditions, with or without liraglutide (20 nM or 100 nM working concentration, in 0.1% DMSO; Selleckchem S8256). Supernatant was collected after stimulation, and islet hormone content was extracted overnight in acid-ethanol (1.5% 12 N HCl in 70% ethanol). Human and mouse insulin levels were quantified by enzyme-linked immunosorbent assay (ELISA) using species-specific commercial kits (Crystal Chem, #90095, 62100), with secretion data normalized to total protein levels for human islets and total hormone content for mouse islets and expressed as percentage release.

### Dynamic secretion assay

Perifusion experiments were performed using the Biorep Perifusion System v4 according to manufacturer specifications. Approximately 50 size-matched islets per chamber were perfused at 100 μL/min with effluent collected at 1 min intervals throughout sequential glucose transitions (2 → 16 → 2 mM. Liraglutide was introduced during the 16 mM glucose phase, followed by 30 mM KCl depolarization during the final low glucose phase. Effluent insulin was measured by ELISA, with results normalized to total DNA content quantified using the PicoGreen dsDNA assay (Life Technologies, P7589).

### Islet cAMP imaging

Islet cAMP dynamics were monitored using the Epac^SH187^–cAMP FRET biosensor (gift from the Piston Lab, Washington University^42,43^). For sensor delivery, intact islets were transduced with adenoviral particles (1×10^11^ particles/mL) in culture medium overnight at 37°C (5% CO_2_), followed by a 48-hour expression period. Prior to imaging, 8-10 islets were plated in rh-laminin-coated (0.7 μg/cm^2^, Gibco) 4-chamber glass-bottom dishes (Cellvis) and equilibrated for 20 min in 11 mM glucose KRBH on the microscope stage.

FRET-based cAMP imaging was performed on a Zeiss LSM 880 confocal microscope using 405 nm excitation and spectral detection across 445-650 nm. Acceptor (mVenus) and donor (mTurquoise) emission signals were separated through linear unmixing (Zeiss ZEN software). Images were acquired at 2.5-second intervals over a 30-minute period. Following an 8-minute baseline recording in 11 mM glucose, liraglutide was added to a final concentration of 100 nM (dissolved in 0.1% DMSO). This concentration was chosen to maximize cAMP production after a prior dose response analysis (20-100 nM) in WT islets. Data analysis was performed by calculating the FRET ratio as the background-subtracted acceptor intensity divided by the background-subtracted donor intensity. For each individual islet, the entire trace was normalized to its own average pre-stimulation baseline.

### Islet Ca^2+^ imaging

Wildtype β-cell GcaMP6f islets were washed in KRBH without glucose, transferred to a laminin-coated 4-chamber glass-bottom dish, starved for 1 hour in KRBH at 3 mM glucose, and then mounted on a climate-controlled stage maintained at 37°C and 5% CO_2_. Ca^2+^ imaging was performed on a Zeiss LSM880 inverted confocal system using a Plan-Apochromat 20×/0.8 M27 oil immersion objective. Images were captured at a frame size of 512 x 512 (708.49 x 708.49 μm^2^) and acquired at a frame time of 471.86 ms. Baseline recordings were performed at 3 mM glucose for approximately 16 min (2,000 cycles), then glucose was raised to 9 mM glucose, and islets were continuously recorded for 39 min (5,000 cycles) to observe first- and second-phase oscillations, then treated with 20 nM liraglutide (in 0.1% DMSO) to mirror the concentration used in the dynamic secretion assay and recorded for an additional 5,000 cycles. For analysis, whole-islet regions of interest (ROIs) were defined, and integrated green fluorescence intensity was used to generate calcium traces in ImageJ. Traces were concatenated and normalized to baseline 3 mM glucose condition. Oscillation parameters in the presence of liraglutide, including peak amplitude, period, active phase (time Ca^2+^ remains above 50% peak amplitude), silent phase (period minus active duration), were quantified using a custom Matlab script (Merrins Lab, https://github.com/hrfoster/Merrins-Lab-Matlab-Scripts)^44^.

### Adenoviral shRNA knockdown

IFT88 knockdown in human islets: Healthy non-diabetic human islets were transduced with adenoviral vectors encoding either GFP-tagged shRNA targeting human *IFT88* (Ad-GFP-h-IFT88-shRNA) or scrambled control (Ad-GFP-U6-scrmb-shRNA, #1122N; Vector Biolabs). Approximately 50 islets per condition were incubated overnight in a low cell attachment plate with viral particles at 2 × 10^9^ IFU/mL concentration. The following day, media was replaced, and islets were maintained for 96 hours with media changes every 48 hours prior to functional assays.

Tulp3 knockdown in mouse islets: Wildtype mouse islets were transduced with adenoviruses expressing Cre recombinase and one of two distinct shRNAs targeting Tulp3 (pAV[Exp]-U6>mTulp3[shRNA#1]-Ins2>Cre and pAV[Exp]-U6>mTulp3[shRNA#2]-Ins2>Cre; VectorBuilder). A stuffer sequence vector (pAV[Exp]-U6>ORF_Stuffer; VectorBuilder) served as control. For each condition, a minimum of 50 islets per mouse were partially dissociated by a 2-minute incubation in Accutase to facilitate viral entry. Islets were transduced (1 × 10^9^ IFU/mL) by co-incubation with virus for 24 hours at 37°C, followed by a 72-hour recovery period in 35 mm dishes with media changes every other day.

### Quantitative PCR

Total mRNA was extracted from transduced islets using RA1 lysis buffer (Macherey-Nagel, 740961) containing β-mercaptoethanol and purified with a Nucleospin RNA kit (Macherey-Nagel, 740955.50). cDNA was synthesized from the purified RNA using a High-Capacity cDNA reverse transcription kit (Thermo Fisher Scientific, 4368814). Quantitative PCR was performed in technical triplicates (QuantStudio 3, Thermo Fisher Scientific, A28131) using Power SYBR Green Master Mix (Thermo Fisher Scientific, 4367659). Relative gene expression was calculated using the 2−ΔΔCt method, with *Gapdh* serving as the endogenous reference gene. Primer sequences are listed in **Table S2**.

### Statistical analysis

Data were analyzed with Microsoft Excel, Image J, MATLAB (for calcium imaging analysis), and GraphPad Prism. All results are presented as mean ± SEM. The specific statistical tests and significance thresholds are detailed in the respective figure legends.

## Acknowledgements

Funding for this study provided by NIH grants R01DK138974 and R01DK140365 to JWH, R01 DK123075, DK125353, DK046492 to JC, K12DK133995 and K08DK142012 to SD, and F30DK143705 to IM. We thank Dr. David Piston for co-mentoring IM and Drs. David Jacobson, Moe Mahjoub, and Thomas Baranski for their valuable guidance as thesis committee members. We are grateful to the Piston Lab for hosting IM and providing critical technical support and advice during the latter phase of this work following the primary PI’s relocation. Human pancreatic islets were provided by the NIDDK-funded Integrated Islet Distribution Program (IIDP) (RRID:SCR_014387) at City of Hope, NIH grant #2UC4DK098085 and the JDRF-funded IIDP Islet Award Initiative. Microscopy was performed at Washington University Center for Cellular Imaging (WUCCI) supported by Washington University School of Medicine, The Children’s Discovery Institute of Washington University and St. Louis Children’s Hospital (CDI-CORE-2015-505 and CDI-CORE-2019-813) and the Foundation for Barnes-Jewish Hospital (3770 and 4642). We thank Dr. Greg Strout (WUCCI) for his expert assistance with immuno-SEM, and Dr. Mark Huising and his laboratory for replicating GLP-1R staining results.

---

**Supplemental Figure S1:**
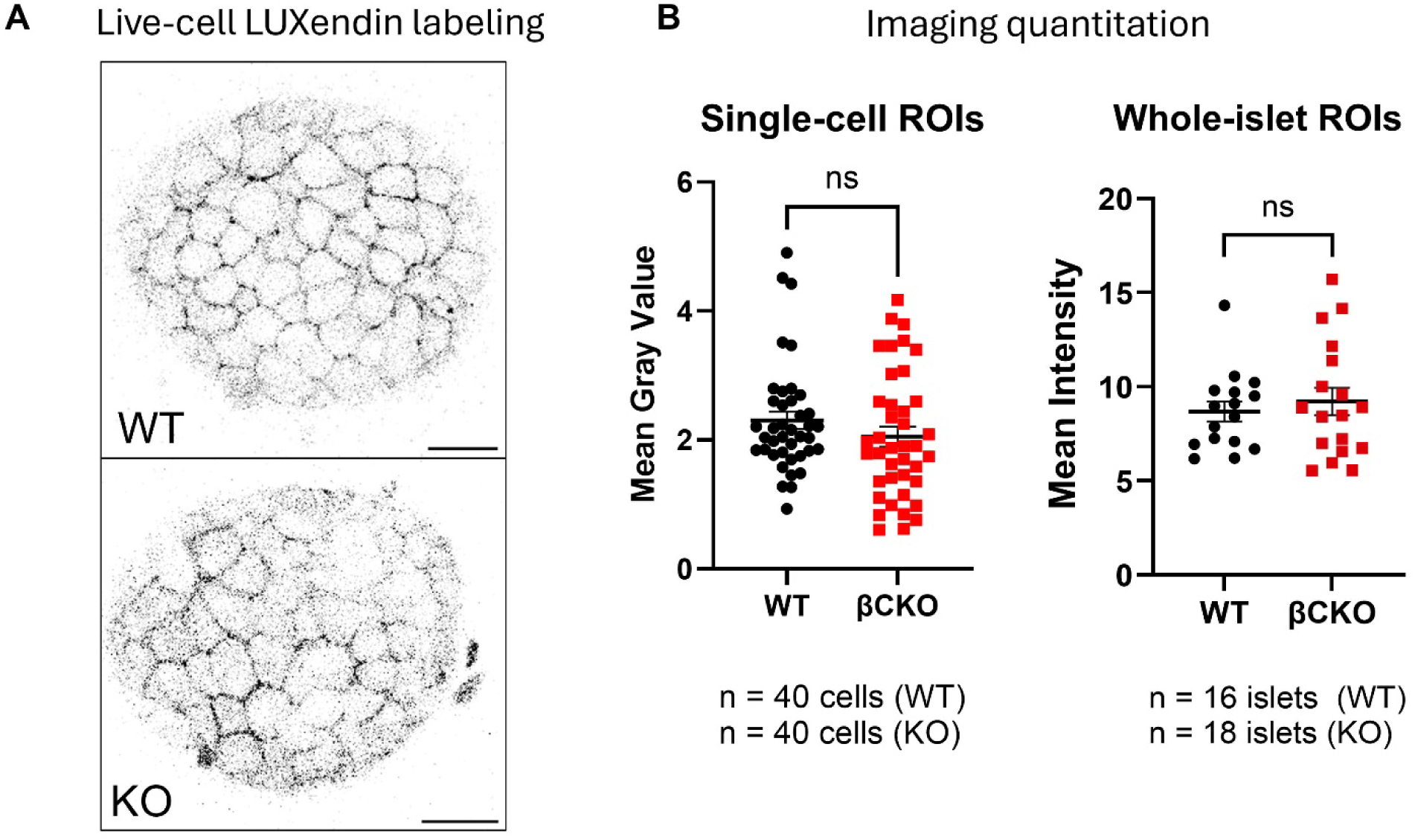
GLP-1R surface expression is unchanged in cilia-deficient islets. (**A**) Live-cell labeling of WT and βCKO islets with the fluorescent GLP-1R antagonist LUXendin645 (100 nM) for 1 hour. (**B**) Quantification of LUXendin645 fluorescence intensities, showing comparable whole-cell/plasma membrane receptor expression. N= 40 single-cell ROIs and 16-18 islets (WT, βCKO). Data presented as mean ±SEM; ns, not significant as assessed by student’s unpaired t-test.

**Supplemental Figure S2:**
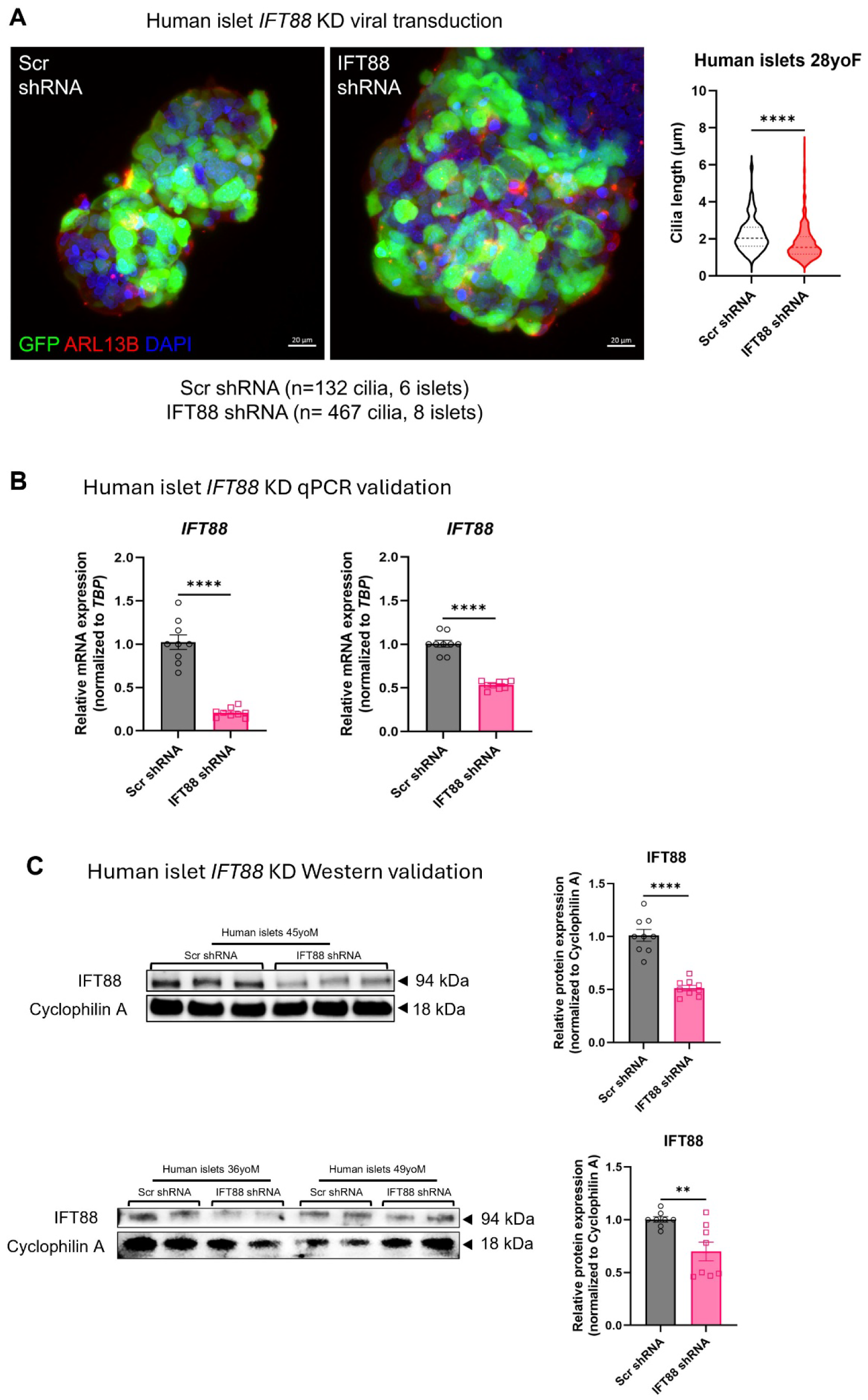
Validation of IFT88 knockdown in human islets. (**A**) Representative z-projected confocal images of human islets (28-year-old female donor) transduced with scrambled (Scr) or *IFT88*-targeting shRNA adenovirus. GFP (green) marks transduced cells; ARL13B (red) labels primary cilia; DAPI (blue) stains nuclei. Scale bar = 20 µm. Cilia length was quantified from both conditions, showing significantly reduced mean length following IFT88 knockdown. n=132 cilia from 6 islets (Scr shRNA) and 467 cilia from 8 islets (IFT88 shRNA). (**B–C**) *IFT88* knockdown efficiency was assessed across multiple donors to demonstrate reproducibility. (**B**) qPCR analysis of IFT88 mRNA levels in islets from two male donors (ages 27 and 44). (**C**) Immunoblot analysis of IFT88 protein levels in islets from three male donors (ages 36, 49 and 45), with corresponding quantifications. Bars represent scrambled control (gray) and *IFT88* shRNA (pink). Data are mean ± SEM of duplicate or triplicate samples per donor. **p < 0.01, ****p < 0.0001, ns = not significant, by student’s unpaired t-test.

**Supplemental Figure S3:**
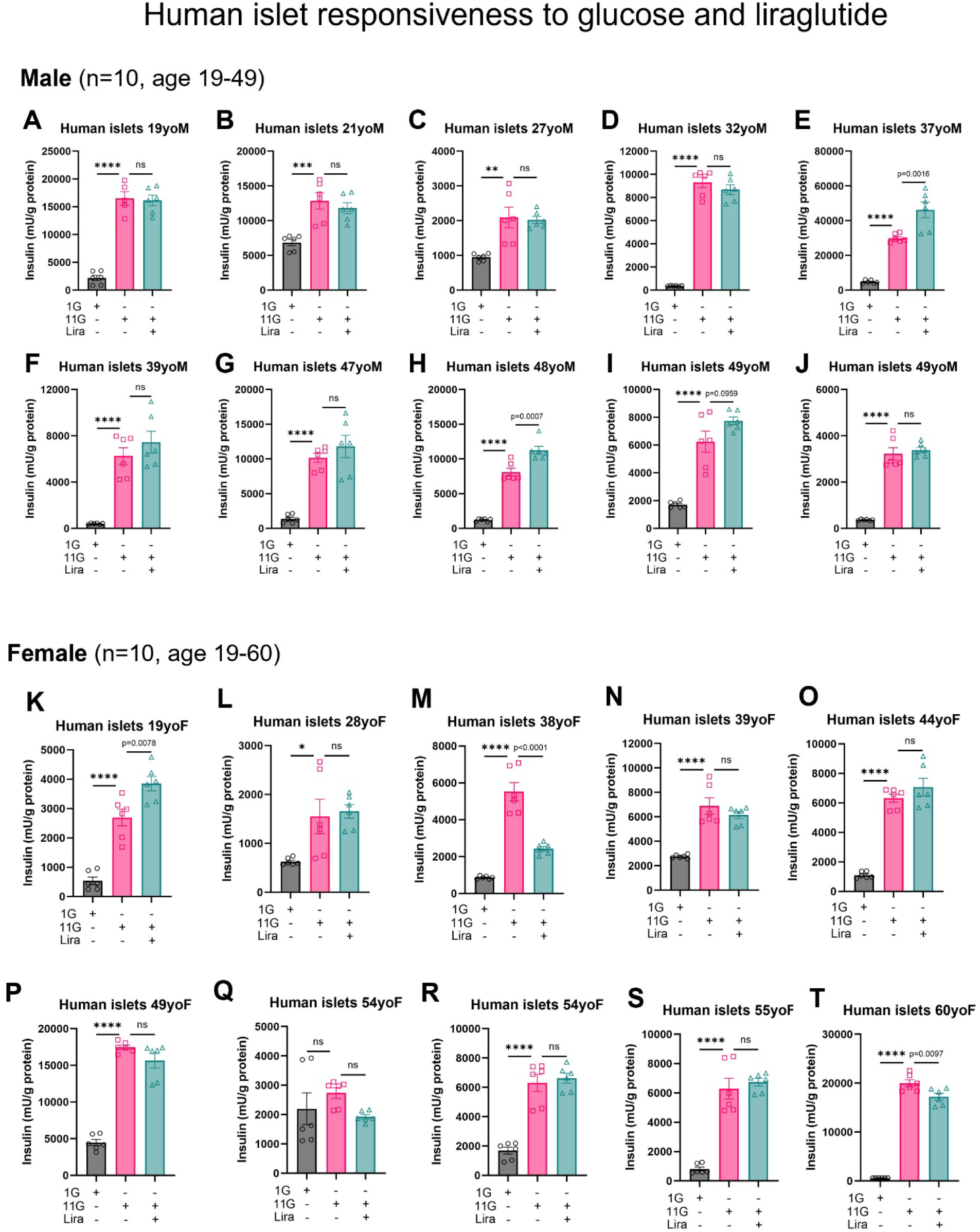
Donor heterogeneity in glucose and liraglutide responsiveness of human islets. (**A-J**) Glucose-stimulated insulin secretion from islets of male donors (ages 19, 21, 27, 32, 37, 39, 47, 48, 49, and 49 years). Secretion was measured at 1 mM glucose (1G), 11 mM glucose (11G), and 11G + 100 nM liraglutide (Lira). (**K–T**) GSIS from islets of female donors (ages 19, 28, 38, 39, 44, 49, 54, 54, 55, and 60 years) under identical conditions. Data are mean ± SEM of triplicate samples per donor. Statistical significance was assessed by one-way ANOVA with Tukey’s multiple comparisons test: *p<0.05, **p<0.01, ***p<0.001, ****p<0.0001; ns, not significant. The variable potentiation by liraglutide across individuals reflects well-documented donor-to-donor heterogeneity in human islet incretin sensitivity.

**Supplemental Figure S4.**
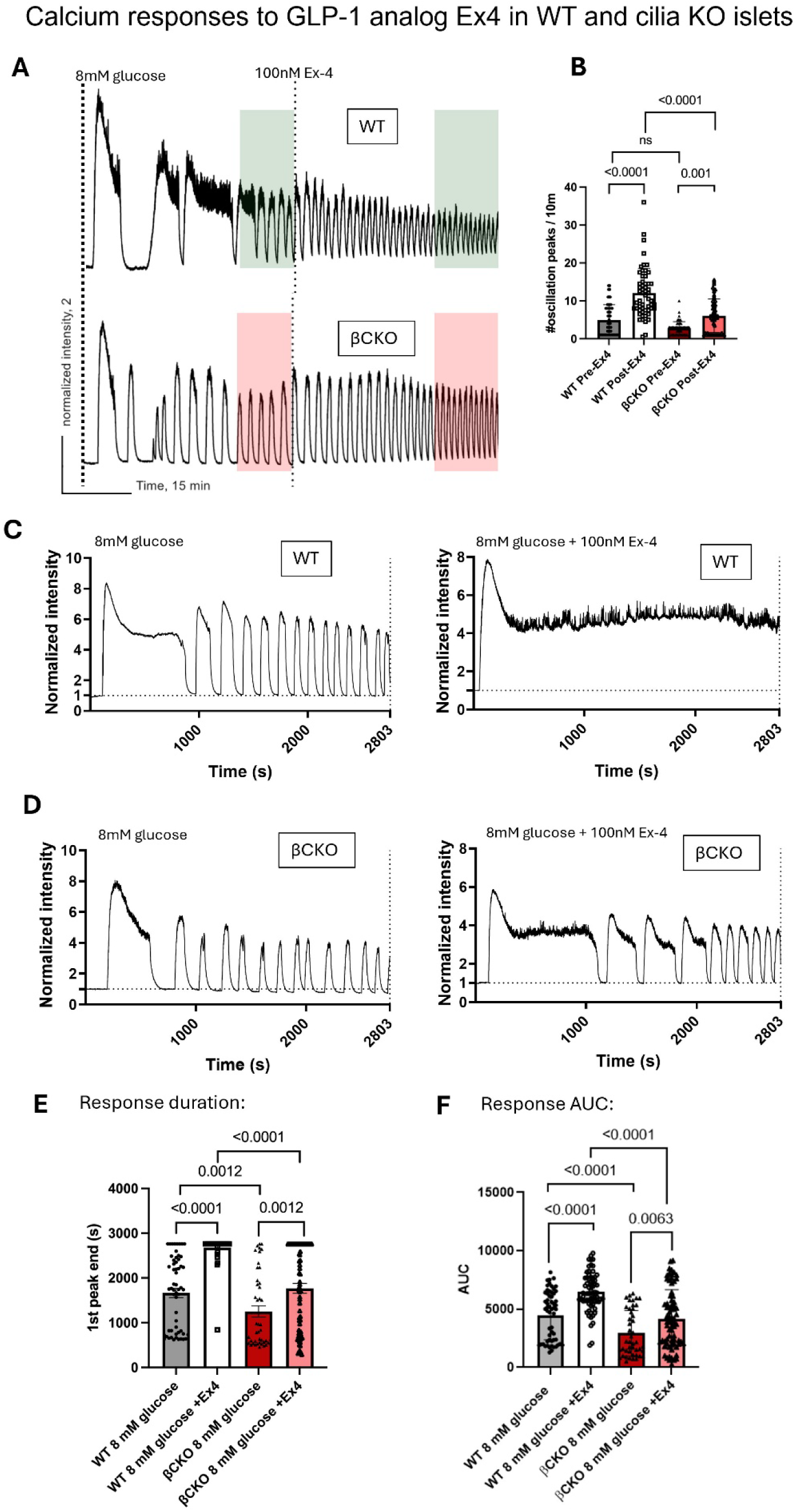
Cilia loss blunts β-cell Ca²⁺ responses to exendin-4, consistent with liraglutide effect. (**A**) Representative live-cell Ca²⁺ (GcaMP6f) recordings from WT (black) and βCKO (red) islets, showing normalized reporter fluorescence over time. Islets were stimulated with 8 mM glucose, followed by the addition of 100 nM exendin-4 (Ex4) at the indicated time. Sampling interval = 5 s. (**B**) Oscillation frequency, quantified as the number of Ca²⁺ peaks per 10-minute interval, before and after Ex4 addition. Ex4 increased oscillation frequency in WT but not in βCKO islets. WT n = 56; βCKO n = 55 islets. ***p < 0.001, two-way ANOVA with Šidák’s *post hoc* test. (**C-D**) Representative single-islet Ca²⁺ traces from an acute stimulation assay. Traces show WT and βCKO islets exposed to 8G alone (left) or to 8G + 100 nM Ex4 added at t = 0 (right). Sampling interval = 0.5 s. I Duration of the first Ca²⁺ transient, measured from the initiation of the Ca²⁺ rise to its return to a sustained baseline. Ex4 prolonged the first transient in both genotypes, but βCKO islets exhibited significantly shorter durations than WT. (**F**) Area under the curve (AUC) of the first Ca²⁺ transient. Ex4 increased AUC in both genotypes; βCKO islets displayed a significantly smaller AUC than WT under all conditions. WT (Glucose only: n = 58; Glucose + Ex4: n = 68); βCKO (Glucose only: n = 45; Glucose + Ex4: n = 73). Data are mean ± SEM. Statistical significance was assessed by two-way ANOVA with Tukey’s post hoc test. *p < 0.05, **p < 0.01, ***p < 0.001; ns, not significant.

