## Supplemental Table 1 for "Primary cilia regulate GLP-1 signaling in pancreatic β cells"

**Supplementary Table S1. Human Islet Donor Characteristics**

Human islet preparation checklist used in this study

Adapted from 1) Hart & Powers (2018) Progress, challenges, and suggestions for using human islets to understand islet biology and human diabetes. *Diabetologia* https://doi.org/10.1007/s00125-018-4772-2, 2) Cottle, Gan, Gilroy *et al.* (2021) Structural and functional polarisation of human pancreatic beta cells in islets from organ donors with and without type 2 diabetes. *Diabetologia* https://doi.org/10.1007/s00125-020-05345-8

| Islet preparation | 1 | 2 | 3 | 4 | 5 | 6 | 7 | 8 | 9 |
|---|---|---|---|---|---|---|---|---|---|
| MANDATORY INFORMATION | | | | | | | | | |
| Unique identifier | SAMN39671770 | SAMN43583143 | SAMN39708909 | SAMN41393792 | SAMN41611427 | SAMN39523303 | SAMN42008301 | SAMN41042507 | SAMN52216388 |
| Donor age (years) | 19 | 19 | 21 | 27 | 28 | 32 | 36 | 37 | 38 |
| Donor sex (M/F) | M | F | M | M | F | M | M | M | F |
| Donor BMI (kg/m$^2$) | 31.7 | 22.3 | 38.1 | 30.1 | 35.0 | 31.3 | 30.0 | 31.3 | 32.4 |
| Donor HbA1c (%) or other measure of blood glucose control (mg/dL) | HbA1c 5.00/ BGL 125.20 | HbA1c 6.00/ BGL 263.40 | HbA1c 5.00/ BGL 157.00 | HbA1c 5.50/ BGL 119.00 | HbA1c 5.80/ BGL 164.60 | HbA1c 5.40/ BGL 207.60 | HbA1c 5.00/ BGL 96.60 | HbA1c 5.60/ BGL 139.20 | HbA1c 5.80/ BGL 228.80 |
| Origin/source of islets | Pennsylvania | Miami | Miami | NCHIBC | NCHIBC | SC-ICRC | Pennsylvania | Pennsylvania | SC-ICRC |
| Islet isolation center | Pennsylvania | Miami | Miami | NCHIBC | NCHIBC | SC-ICRC | Pennsylvania | Pennsylvania | SC-ICRC |
| Donor history of diabetes? Please | No | No | No | No | No | No | No | No | No |

| | | | | | | | | | |
|---|---|---|---|---|---|---|---|---|---|
| select yes/no from drop down list | | | | | | | | | |
| If Yes, complete the next two lines if this information is available | | | | | | | | | |
| Diabetes duration (years) | | | | | | | | | |
| Glucose-lowering therapy at time of death | | | | | | | | | |
| RECOMMENDED INFORMATION | | | | | | | | | |
| Donor cause of death | Head trauma | Anoxia | Head trauma | Anoxia | Anoxia | Anoxia | Anoxia | Anoxia | Cerebro vascular /stroke |
| Warm ischaemia time (h) | No | No | No | No | No | Yes | No | No | No |
| Cold ischaemia time (hh:mm) | 18:20 | 05:20 | 16:21 | 08:30 | 06:55 | 08:57 | 14:07 | 09:25 | 05:32 |
| Estimated purity (%) | 85 | 85 | 95 | 90 | 80 | 78 | 80 | 90 | 80 |
| Estimated viability (%) | 89 | 93 | 90 | 85 | 85 | 95 | 97 | 96 | 95 |
| Total culture time (h) | 71 | 95 | 63 | 25 | 173 | 163 | 119 | 161 | 56 |
| Glucose-stimulated insulin secretion or other functional measurement | 4.3 | 2.9 | 5.1 | 5.0 | 2.0 | 1.9 | 1.3 | 2.0 | NA |
| Handpicked to purity? Please select yes/no from drop down list | No | No | No | No | No | No | No | No | No |
| Additional notes | NA | NA | NA | NA | NA | NA | NA | NA | NA |

| Islet preparation | 10 | 11 | 12 | 13 | 14 | 15 | 16 | 17 | 18 |
|---|---|---|---|---|---|---|---|---|---|
| MANDATORY INFORMATION | | | | | | | | | |
| Unique identifier | SAMN39912961 | SAMN41457782 | SAMN42742099 | SAMN51198036 | SAMN46865134 | SAMN40380406 | SAMN40963318 | SAMN39447621 | SAMN39639106 |
| Donor age (years) | 39 | 39 | 44 | 44 | 45 | 47 | 48 | 49 | 49 |
| Donor sex (M/F) | M | F | M | F | M | M | M | M | M |
| Donor BMI (kg/m$^2$) | 31.0 | 37.1 | 33.1 | 31.0 | 26.6 | 34.6 | 36.5 | 34.4 | 32.5 |
| Donor HbA1c (%) or other measure of blood glucose control (mg/dL) | HbA1c 5.80/ BGL 188.20 | HbA1c 5.70/ BGL 163.60 | HbA1c 5.90/ BGL 139.00 | HbA1c 4.50/ BGL 148.40 | HbA1c 5.70/ BGL 186.40 | HbA1c 5.50/ BGL 150.80 | HbA1c 5.60/ BGL 151.20 | HbA1c 5.50/ BGL 165.20 | HbA1c 5.00/ BGL 160.40 |
| Origin/source of islets | Wisconsin | Pennsylvania | Prodo Labs | Arizona | Pennsylvania | Prodo Labs | Wisconsin | SC-ICRC | Imagine |
| Islet isolation center | Wisconsin | Pennsylvania | Prodo Labs | Arizona | Pennsylvania | Prodo Labs | Wisconsin | SC-ICRC | Imagine |
| Donor history of diabetes? Please select yes/no from drop down list | No | No | No | No | No | No | No | No | No |
| If yes, complete the next two lines if this information is available | | | | | | | | | |
| Diabetes duration (years) | | | | | | | | | |

| | | | | | | | | | |
|---|---|---|---|---|---|---|---|---|---|
| Glucose-lowering therapy at time of death | | | | | | | | | |
| RECOMMENDED INFORMATION | | | | | | | | | |
| Donor cause of death | Head trauma | Cerebro vascular /stroke | Cerebro vascular /stroke | Cerebro vascular /stroke | Head trauma | Cerebro vascular /stroke | Anoxia | Anoxia | Cerebro vascular /stroke |
| Warm ischaemia time (h) | Yes | No | No | No | No | No | Yes | Yes | No |
| Cold ischaemia time (hh:mm) | 07:30 | 10:19 | 10:45 | 7:56 | 04:41 | 08:05 | 09:00 | 11:37 | 04:36 |
| Estimated purity (%) | 90 | 90 | 90 | 90 | 90 | 90 | 95 | 95 | 85 |
| Estimated viability (%) | 98 | 88 | 95 | 97 | 94 | 95 | 98 | 95 | 89 |
| Total culture time (h) | 89 | 164 | 106 | 61 | 200 | 158 | 107 | 72 | 97 |
| Glucose-stimulated insulin secretion or other functional measurement | 1.7 | 1.7 | NA | NA | 1.5 | 3.3 | 2.6 | 1.2 | 2.7 |
| Handpicked to purity? Please select yes/no from drop down list | No | No | No | No | No | No | No | No | No |
| Additional notes | NA | NA | NA | NA | NA | NA | NA | NA | NA |

| Islet preparation | 19 | 20 | 21 | 22 | 23 | 24 |
|---|---|---|---|---|---|---|
| MANDATORY INFORMATION | | | | | | |
| Unique identifier | SAMN43823987 | SAMN47406629 | SAMN40867047 | SAMN43463703 | SAMN45232023 | SAMN39980165 |
| Donor age (years) | 49 | 52 | 54 | 54 | 55 | 60 |
| Donor sex (M/F) | F | F | F | F | F | F |
| Donor BMI (kg/m$^2$) | 37.6 | 30.5 | 27.2 | 28.7 | 29.6 | 29.6 |
| Donor HbA1c (%) or other measure of blood glucose control (mg/dL) | HbA1c 5.60/ BGL 138.00 | HbA1c 5.00/ BGL 211.40 | HbA1c 5.00/ BGL 169.00 | HbA1c 5.70/ BGL 121.60 | HbA1c 5.10/ BGL 94.40 | HbA1c 6.20/ BGL 155.60 |
| Origin/source of islets | SC-ICRC | Imagine | NCHIBC | Prodo Labs | Prodo Labs | SC-ICRC |
| Islet isolation center | SC-ICRC | Imagine | NCHIBC | Prodo Labs | Prodo Labs | SC-ICRC |
| Donor history of diabetes? Please select yes/no from drop down list | No | No | No | No | No | No |
| If yes, complete the next two lines if this information is available | | | | | | |
| Diabetes duration (years) | | | | | | |

| | | | | | | |
|---|---|---|---|---|---|---|
| Glucose-lowering therapy at time of death | | | | | | |
| RECOMMENDED INFORMATION | | | | | | |
| Donor cause of death | Cerebro vascular /stroke | Head trauma | Anoxia | Head trauma | Cerebro vascular /stroke | Cerebro vascular /stroke |
| Warm ischaemia time (h) | No | No | NA | No | No | Yes |
| Cold ischaemia time (hh:mm) | 08:37 | 10:26 | NA | 07:02 | 08:10 | 09:17 |
| Estimated purity (%) | 90 | 85 | 90 | 90 | 85 | 90 |
| Estimated viability (%) | 95 | 92 | 85 | 95 | 95 | 95 |
| Total culture time (h) | 73 | 258 | NA | 240 | 209 | 159 |
| Glucose-stimulated insulin secretion or other functional measurement | 2.8 | 4.7 | NA | NA | NA | 1.8 |
| Handpicked to purity? Please select yes/no from drop down list | No | No | No | No | No | No |
| Additional notes | NA | NA | NA | NA | NA | NA |
