## Supplemental Table 2 for "Primary cilia regulate GLP-1 signaling in pancreatic β cells"

**Supplementary Table S2**

**Primer sequences for quantitative RT-PCR**

| Target gene | | Primer sequence |
|---|---|---|
| *Glp1r* | Forward | CGGAGTGTGAAGAGTCTAAGCG |
| | Reverse | ATGGCTGAAGCGATGACCAAGG |
| *Gapdh* | Forward | AGGTTGTCTCCTGCGACTTCA |
| | Reverse | CAGGAAATGAGCTTGACAAAGTTG |
| *Tulp3* | Forward | CAGCTGAAGCTGGACAATCA |
| | Reverse | GGGTTTGGCTGTACCATGAG |
| *IFT88 (human)* | Forward | TCGGCTAGATGAGGCTTTGGAC |
| | Reverse | CACTGACCACCTGCATTAGCCA |
| *TBP (human)* | Forward | GCCATAAGGCATCATTGGAC |
| | Reverse | AACAACAGCCTGCCACCTTA |
